# Neurokraken: A fully flexible, open-source, python-based neuroscience behavior platform

**DOI:** 10.64898/2026.06.30.735592

**Authors:** Alexander Wallerus, Sofia Castro e Almeida, Johannes Passecker

## Abstract

A major challenge in behavioral neuroscience is the lack of a unified software framework capable of implementing diverse paradigms across species and experimental setups. Researchers currently face a trade-off: they must either spend significant time developing custom, siloed solutions that hinder reproducibility, or incur substantial costs purchasing inflexible, closed systems. Here, we present Neurokraken, an open-source, Python-native platform designed to overcome these limitations. Neurokraken allows writing experiment progression entirely in standard python, while its core architecture automatically sets up a microcontroller for the connected hardware components and enables python side access with millisecond-precision timing and automatic logging. The system prioritizes ease of use and flexibility, enabling advanced series of events and conditions, the usage of python ecosystem code and packages within experiments, and the addition of any arduino-compatible electronic devices for custom experiments. As a result, users can easily create interactive virtual and real environments to engage, monitor, and record subjects. We present Neurokraken’s versatility across a wide range of paradigms, for human and non-human primate psychophysics, and complex rodent behavior in both head-fixed and freely moving paradigms. Its modular design allows for rapid hardware reconfiguration, while a fully customizable user interface enables real-time monitoring and interactive experimental control without compromising timing precision. By uniting laboratory-grade precision with an accessible and flexible open-source philosophy, Neurokraken provides a single, powerful solution to design and execute next-generation behavioral experiments. We hope Neurokraken helps accelerate research, improve reproducibility throughout the neuroscience community, and make advanced behavioral experimentation more accessible through its substantial cost-efficiency.

## Introduction

Behavior is the primary output of the nervous system and the ultimate benchmark for understanding its function. For decades, behavioral neuroscience has relied on well-established, often simplified paradigms to dissect the neural basis of learning, perception, and action. While these approaches have been invaluable, the field faces a growing need for computational neuroethological paradigms that preserve experimental control while better aligning with the richness of real-world cognition^1^. Alongside, neuroscience is undergoing a transformation, fueled by two parallel advances. First, new techniques for recording and manipulating neural activity provide unprecedented access to the brain in action^2,3,3–6^. Second, the rise of machine learning and computational theory offers powerful, species-agnostic frameworks for interpreting complex neural and behavioral data^1,7–11^. Together, these advances also necessitate new behavioral paradigms, ones that are sufficiently complex and/or naturalistic to test new hypotheses.

This need has spurred the development of several new experimental platforms over the last decades^12–18^. Commercial “turn-key” systems offer reliability but are often prohibitively expensive, mostly with closed-source architectures that limit customization. In parallel, a vibrant ecosystem of open-source platforms has emerged, with numerous tools becoming cornerstones for many labs interested in rodent, non-human primate and human behavior and its psychophysics, respectively^1,12,19–25^. While these community-driven efforts have been instrumental for many research labs across the globe, critical gaps remain. Most platforms remain optimized for a specific task modality, limiting the type and number of devices that can be connected and are limited to a specific species. This increases the cost and time effort for adapting, translating and creating new tasks. Particularly the integration of rich visual and virtual environments, comparing behavior to artificial agents, or deploying real-time computer vision often requires extensive customization, fragmenting the experimental workflow and compromising reproducibility. This often creates “island” solutions across and within research labs that hinder comparability, translatability, and reproducibility. Consequently, the field lacks single frameworks that are powerful enough for complex paradigms, yet flexible enough to bridge species and experimental setups, all while remaining fully open and accessible.

To bridge this gap, we developed Neurokraken: an open-source, Python-native platform for designing and executing next-generation behavioral experiments. Its core principle is the clean separation of experimental logic from hardware implementation. This allows researchers to define complex tasks using intuitive Python code, while a dedicated microcontroller backend with an automated configuration workflow ensures millisecond-precision timing and reliable device control. The framework is designed for straightforward integration with diverse hardware arrangements, from simple sensors to advanced commercial optogenetic and electrophysiology systems. Critically, it can help unify research across domains by enabling the same task logic to be deployed in rodents, non-human primates, humans, and even artificial agents. By uniting laboratory-grade precision with the flexibility of an open-source, Python-native philosophy, Neurokraken provides a solution designed to accelerate progress in behavioral and systems neuroscience in an open-source manner for increased inclusivity given its high cost-reduction capabilities. By removing proprietary barriers and relying on affordable, off-the-shelf components, Neurokraken fosters global scientific equity, democratizing access to cutting-edge behavioral research for underfunded institutions. The presented paradigms achieve high precision and full customizability for only a few hundred euros, leveraging the flexible integration of affordable, reliable off-the-shelf hardware and 3D-printed components.

## Results

### A Unified Framework for Cross-Species Behavioral Neuroscience

Neurokraken provides a single, Python-native framework with the capacity for implementing an array of behavioral tasks across species, sensory modalities, and experimental goals in a fully open-source and cost-effective manner. The platform was designed to enable easy expansion for established labs due to a wide integrability of devices without compromises on precision, while enabling easy starts for individuals new to behavioral task-designs. To exemplify the flexibility of Neurokraken we highlight several examples across species, aligned with simple documentation to enable easy task development for researchers across fields (Fig. 1, see https://neurokraken.github.io/Neurokraken-book). Neurokraken’s versatility also allows for direct comparisons between biological and artificial intelligence. For instance, Neurokraken tasks can use python packages like ViZDoom, a reinforcement-learning environment, with sensor/input device or agent actions and retrieved visuals, enabling the same task to be performed by an artificial agent (Fig. 1a) or a living organism. This versatility extends to simple or complex spatial navigation tasks, from go/no-go exploration in self-designable virtual corridors (Fig. 1b) to 2D paradigms where in our example head-fixed mice use a joystick for navigation (Fig. 1e). The framework supports an open range of input devices and sensors, from touch, wheel rotation, and light beam break to joysticks, specialized research equipment and standard keyboards depending on the needs of the experiment (Fig. 1e, 1f and 1h) with extendibility as a focus point. This also applies to task-controlled devices from lights, valves and motor-movable elements to sounds and visuals on displays. For human psychophysics, Neurokraken supports classic paradigms requiring high temporal precision and real-time monitoring. For example, Neurokraken can readily accommodate perceptual decision-making tasks such as the random-dot motion discrimination task (Fig. 1c), where participants indicate choices via touchscreen input. Neurokraken integrates cameras to record and provide frames for live-visualization and online-analysis. Coupled with Python ecosystem packages, this allows for easy eye tracking for studying visual targeting and saccades (Fig. 1d). In rodent research, the framework accommodates both head-fixed and freely moving paradigms (Fig. 1e-h). Experiments can be augmented with measurements and analysis from computer vision, ML, and AI, such as continuous real-time tracking of pupils or movement trajectories, utilizing the video object segmentation tool CUTIE^26^ (Fig. 1f and 1g) or other tracking approaches as needed. These examples highlight Neurokraken’s capacity to serve as a comprehensive platform for modern behavioral neuroscience.

**Figure 1:**
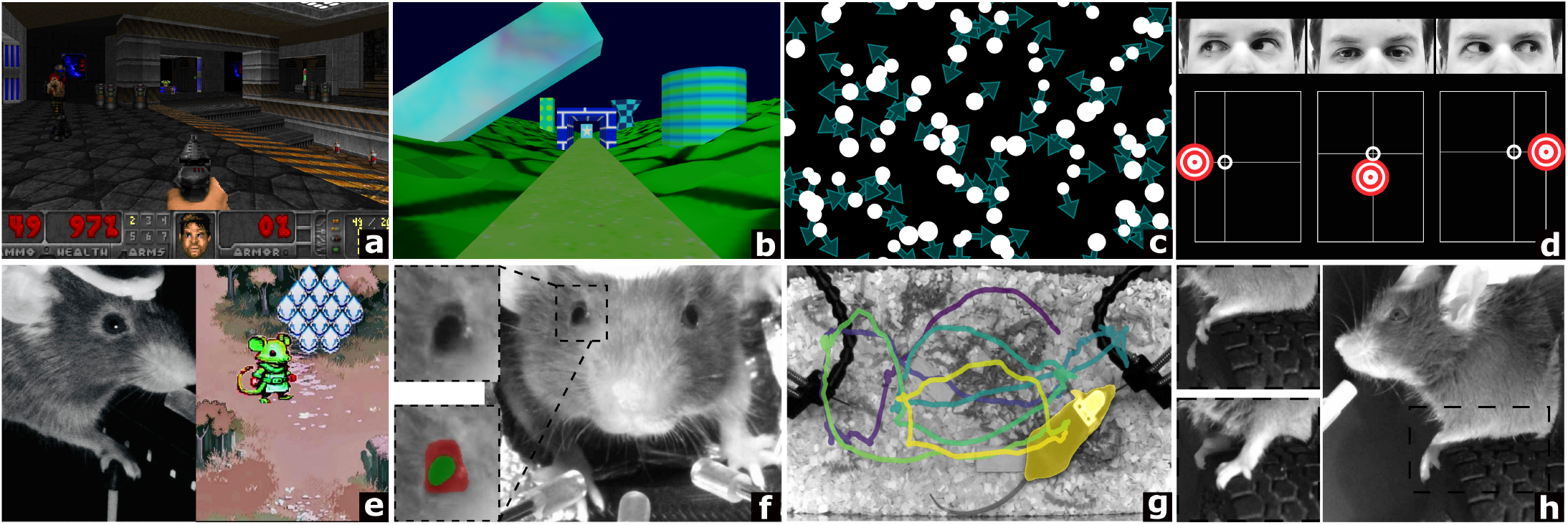
Neurokraken enables flexible behavioral paradigms across a wide range of environments, subjects and neuroscience applications. Task examples include **a):** integration of a python AI reinforcement learning environment (ViZDoom) **b):** a traversable customized 3D virtual corridor for go/no-go spatial learning **c):** a random dot motion task in which participants use a touchscreen to indicate the perceived dominant direction of motion in a dot field **d):** eye-tracking integration for visual targeting in a human saccade task **e):** a head-fixed mouse navigates a virtual environment using a joystick to collect rewards **f):** a mouse performs an olfaction-based three-choice decision-making task with real-time tracking of pupil dynamics **g):** a mouse alternates between two reward spouts while its position is tracked in real time - colored traces indicate previous walking paths **h):** a head-fixed mouse performing a wheel-based decision-making task, steering a displayed shape left or right to receive a reward.

#### Visual versatility

To support users, multiple simulation modes allow tasks to be developed, tested, and debugged without connected task hardware but via keyboard-driven user inputs or scripted agents -accelerating iteration and facilitating collaboration between theorists and experimentalists. Documentation, exemplars, and step-by-step guides support onboarding (see methods and online documentation for more details). For example, the documentation helps users to readily create with the py5 library, fully compatible with Neurokraken’s Python-native architecture, rich visual and interactive environments. Py5 (https://py5coding.org) is a Python version of the well-established *Processing* (https://github.com/processing creative coding environment. Thus, 2D text and shapes (Fig. 1c and 1d) as well as self -made or retrieved images and textures (Fig. 1e and Fig. 1a) can easily be placed, timed, and animated according to task- and sensor data across one or more displays. This also applies to 3D models that can be created in software like blender (https://blender.org) as well as the location of the 3D-scene-camera which enables rendering rich and highly custom task visuals like moving through a 3D environment with a specific position or landmark rewarding movement stops (Fig. 1b)

### Flexible Architecture for Rapid and Reproducible Development

Neurokraken is a modular platform combining a Python-based control layer with a microcontroller for precise timing and synchronization. Python specifies the task logic, while the Teensy handles millisecond-resolution event timing, coordinating sensor sampling and actuator control, such as triggering connected valves. Central to Neurokraken’s adaptability is its software architecture where experiments are defined entirely in Python, using intuitive constructs like conditionals and function calls (Fig. 2, left panel), removing the limitations of specialized scripting languages and experimental configuration files. This architecture is built on two core layers: i) A setup layer defines the experimental hardware (e.g., sensors, stimuli, displays, cameras, microphones), allowing a task’s code to run flexibly on different rigs and with different types of hardware ii) A logic layer contains the behavioral paradigm’s progression, such as stimulus contingencies, trial structures, parameter calculations, and adaptive adjustments. The framework does not impose a rigid structure, empowering researchers to implement tasks, such as continuous control loops, event sequences, or state machines. In general, Neurokraken can be run in two distinct modes to accommodate different laboratory workflows. *Imported Mode*: This standard approach allows a user to import Neurokraken as a package into their Python script. It offers maximum flexibility and is ideal for task development, debugging, and integration into larger, custom pipelines. *Runner Mode*: This command-line utility (*kraken*.*bat*) is designed for high-throughput, standardized experimental sessions that may include numerous tasks and task-variations. It operates on a specified task folder containing separate files for configuration (*config*.*py*), experimental logic (task.*py*), and optionally the UI. The Runner Mode can accept common arguments (e.g., enable keyboard mode, disable logging) and prompt the user for session-specific metadata (e.g., subject ID, weight) before starting the experiment. It promotes consistency, traceability and high-quality data collection, by automatically saving a backup of the correspondent experiment code. This dual-mode system supports both rapid, flexible development and robust, reproducible deployment in a research environment. Further, the dual-mode architecture inherently provides high-throughput scalability required for large-scale, standardized behavioral screening facilities without sacrificing data provenance.

**Figure 2:**
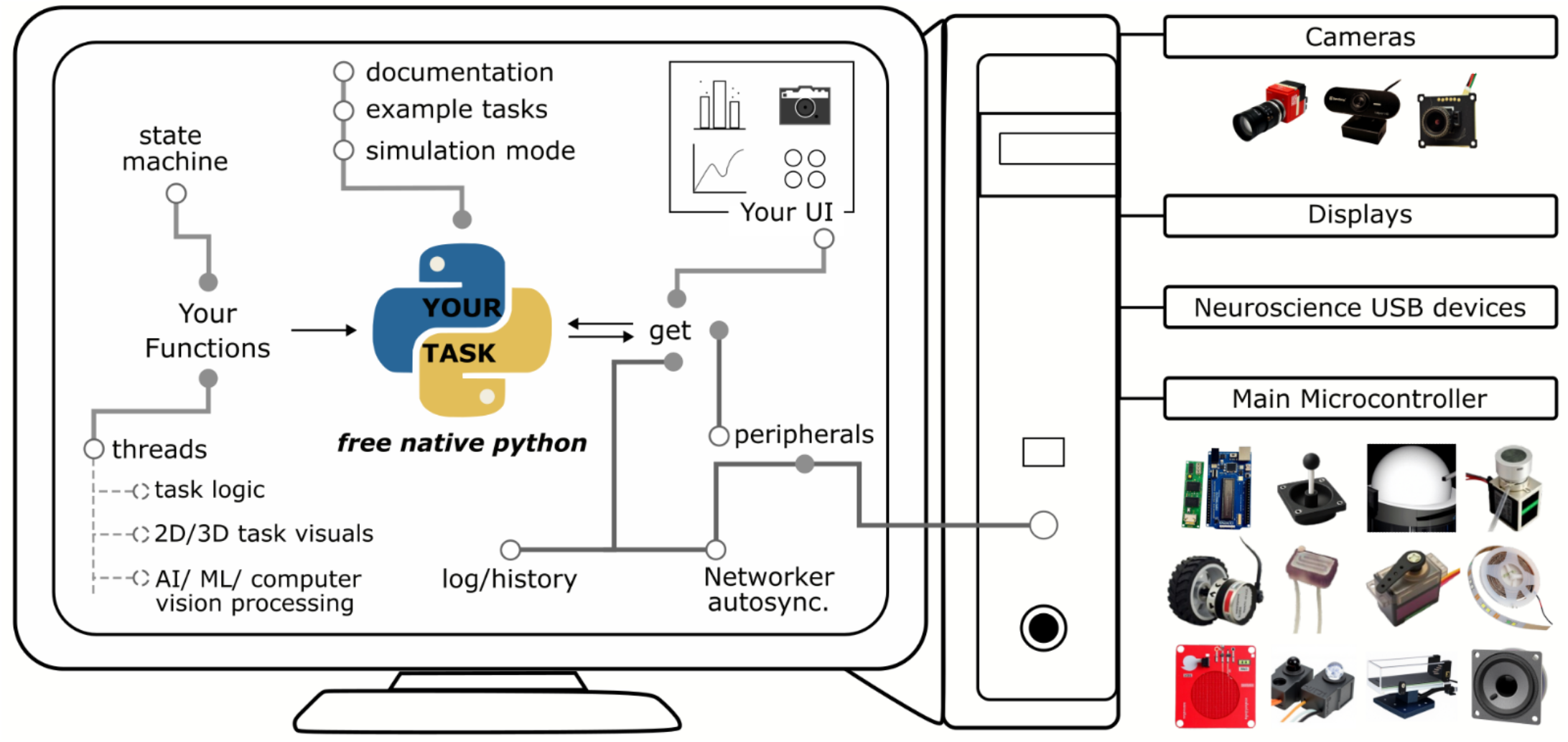
Neurokraken’s Software Architecture and Hardware Integration. Left panel: Python-based Software Environment. The framework leverages a fully Python-native environment, granting users flexibility and access to a rich ecosystem of packages. A central task structure defines the experimental logic, orchestrating all modules - such as the state machine, visual rendering, and user-defined extensions - through intuitive function calls. Computationally intensive processes, like real-time computer vision or AI model inference, operate in parallel processing threads to ensure they do not interfere with the main experiment’s timing. Development is accelerated by comprehensive documentation, a wide range of example tasks, and simulation modes for testing code without connected hardware. Neurokraken’s get serves as the main interface to interact with components like accessing cameras, reading sensors, controlling stimuli, and accessing task variables as well as the automatically logged history of devices. This makes it also useful to interact with the task from outside elements like a UI to display live-data or add overrides. **Right panel: Versatile Hardware Platform**. The software seamlessly interfaces with a diverse array of hardware components. Neurokraken natively supports common peripherals like cameras (from consumer-grade webcams to high-performance GenICAM systems), displays, and microphones. It also integrates and synchronizes with specialized commercial neuroscience equipment, including optogenetic and electrophysiology rigs. At the core of its hardware control is a central microcontroller that manages an extensible suite of devices, such as sensors, motors, valves, lights, and rotary encoders.

#### High-Precision Data Synchronization via the Networker

The system integrates high-performance timing, hardware control, and data logging, through a structured Python API communicating with an Arduino-compatible Teensy 4.1 microcontroller. All communication between the Python script and the microcontroller is managed by a core component called the Networker. To ensure data integrity and timing precision, the Networker maintains a time-stamped list of changes in sensor readings (e.g., licks, movements) with millisecond precision. During each communication, the Networker transmits this complete buffer to the Python side for automatic logging and real-time access. This architecture guarantees that no data points are lost, even if the host computer experiences minor processing delays, thus preserving the temporal fidelity of the behavioral and sensor data. A modifiable pulse clock further enables high-precision synchronization with external devices, often necessary for complex recording setups. Each session is captured in a structured “.json” log file that includes subject/session metadata, as well as timestamped device readouts (e.g., sensor traces, stimulus updates, video frames, audio) and utilized task events (state progression, trials, blocks). The log can be dynamically extended by researchers as part of the experiment. Furthermore, this centralized, format-agnostic logging approach perfectly positions the output data for straightforward integration into standardized, community-driven data ecosystems like Neurodata Without Borders. Real-time interaction with the Neurokraken’s managed components is mediated through a centralized control object (*get*). This object serves as a simple and unified API to read from sensors, send commands to actuators, log data, and control the flow of the experiment from any part of the code. This access point enables parallel access to experiments from user interfaces, or advanced applications such as running a real-time pose estimation model in a parallel thread that uses *get*.*camera* to access live video frames and trigger events based on an animal’s behavior.

Neurokraken interfaces with a broad range of hardware (see Fig. 2 right panel), including multiple simultaneous cameras (from webcams to advanced GenICAM systems), microphones, displays, speakers, microcontroller-controlled peripherals (sensors, motors, valves, lights), general-purpose USB devices, and neuroscience-specific equipment (e.g., commercial optogenetics and electrophysiology rigs). Critically, this integration is designed for flexibility. The hardware abstraction allows switching from any input (e.g., touch sensor, beam break, or rotary encoder) or output peripherals (e.g., valves, motors) without re-authoring the task logic, enabling rapid adaptation across paradigms. To eliminate the need for manual microcontroller programming, the framework provides a utility script which parses the Python hardware configuration and automatically generates for established and self-added new devices the corresponding C++ header file for the Teensy microcontroller. The researcher only needs to upload the pre-built firmware alongside this generated file, ensuring the hardware setup matches from the Python-side configuration without writing or debugging any C++ code. This automated ‘Arduino-free’ workflow reduces the traditional hardware training bottleneck, preventing skill diversion and allowing neuroscientists to focus on experimental design rather than low-level engineering. The respective device library is designed for easy extensions. To add a new custom device, a user only needs to create a simple Arduino class that inherits from a *Control, Sensor*, or *Process* base class (see online documentation for more details).

To evaluate the stability, speed, and timing precision of Neurokraken under varying computational loads, we conducted performance benchmarks across five distinct conditions. These conditions were designed to reflect a spectrum of realistic real-world experimental use cases, ranging from low-demand setups to high-load scenarios involving high-frame-rate video acquisition, real-time computer vision analysis using the CUTIE vision transformer, and a high density of 32 active inputs and controlled peripherals (Supplementary Table 1). In all test conditions, the Networker communicated changes at highly reliable millisecond precision without loss, even under increasing computational demand (sampling jitter of 0.3-1.08μs with communication interval jitter of 0.48-0.69ms, Supplementary Table 1). We further confirmed the reliability of the pulse clock as a ground truth signal for synchronizing external devices with microsecond precision (jitter of 0.9 μs - 62 μs, depending on load).

### Interactive Experimental Control via a Fully Customizable User Interface

Moving beyond the “black-box” paradigm, the framework can provide live access to all task variables, state progression, performance metrics, task logic, hardware status, and hardware controls. This enables researchers to monitor behavior as it unfolds and, when necessary, intervene dynamically by adjusting parameters, overriding task rules, or issuing manual triggers when necessary. All user actions - whether issued through buttons, toggles, sliders, or text inputs - are instantly executed without interrupting the task’s timing or core logic.

A key feature of Neurokraken’s architecture is that it is UI-agnostic. As access to Neurokraken’s components is facilitated through the general *get* interface, Python-based UI frameworks like *PySide6, Tkinter or KrakenGUI* can be used to create customized real-time observation and control interfaces (Fig. 3a). This design enables users to create task-specific dashboards tailored to their needs, whether for displaying raw video feeds, psychometric functions, device states, or live trial histories with performance analysis.

**Figure 3:**
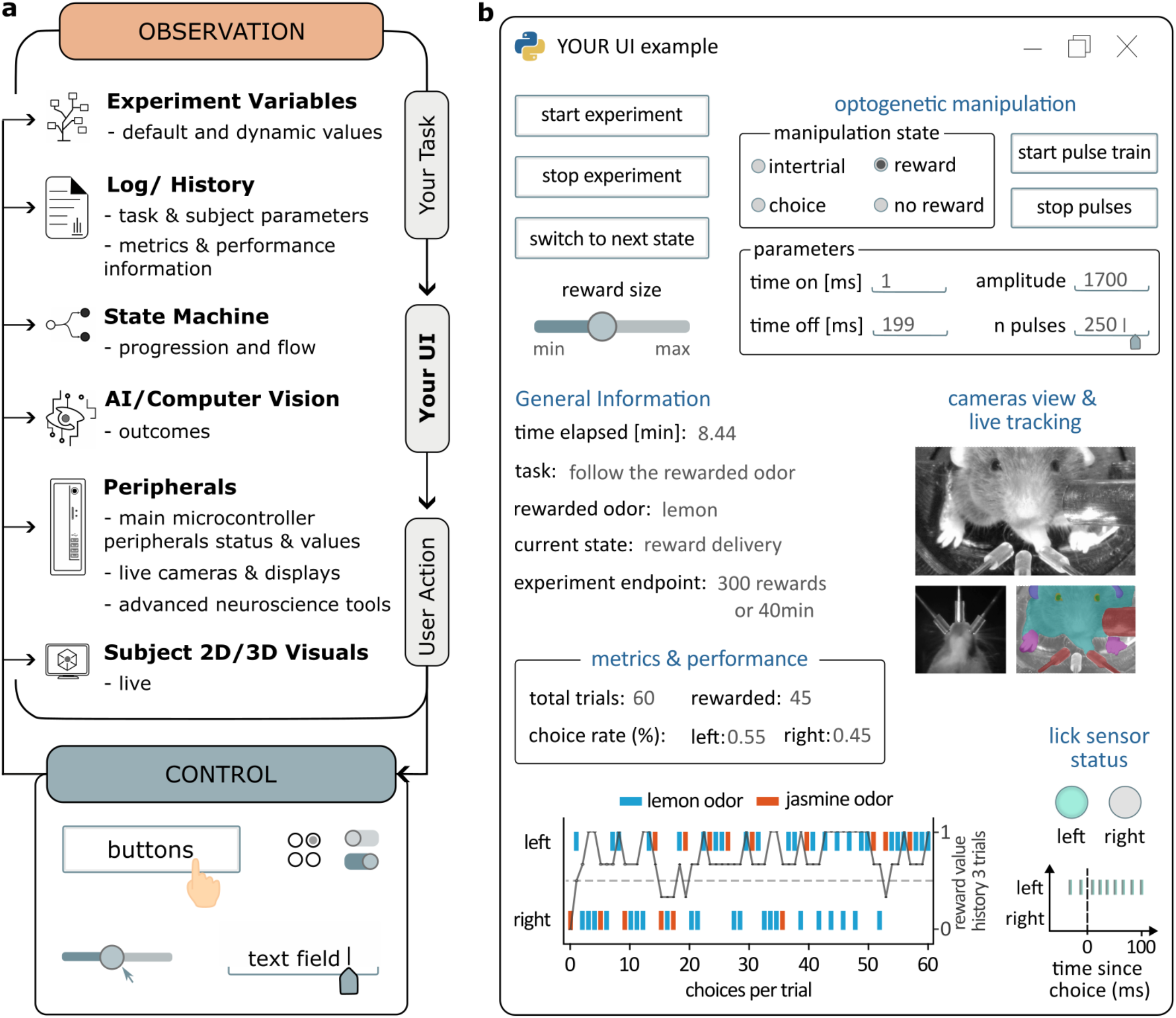
Real-Time Experimental Monitoring and Control via a Flexible UI Architecture. **a)** schematic of the UI Structure. Neurokraken’s open architecture allows researchers to build custom Interfaces using python-based UI frameworks. The UI provides a bidirectional link to the running experiment, enabling two primary functions. i) The observation panel can display all aspects of the task in real time, including experimental variables, state machine progression, live performance analytics, and the status of hardware peripherals. ii) Concurrently, the control panel allows researchers to interact with the task dynamically by adjusting parameters, manually triggering events, triggering changes to actuators or stimuli, overriding the experiment state or simply adding time-stamped notes to the log using elements like buttons, sliders, and text fields depending on the need of the experiment. **b) Example UI during a behavioral task utilizing optogenetics**. This interface demonstrates a practical application of these principles in a rodent experiment. Researchers have full command over the task’s state machine (e.g., start, stop) and fine-grained control over optogenetic stimulation protocol settings, allowing for real-time adjustment of stimulation epochs, pulse parameters, and power. The monitoring section displays key task variables and dynamically plots trial-by-trial performance. Integrated peripheral feedback includes status indicators for lick sensors and multiple live camera views. These video streams can be processed, for example with **Neurokraken’s** CUTIE model utilities, for real-time tracking of the animal’s movement and posture

To illustrate these principles, Figure 3b shows an example interface designed for an optogenetics-based behavioral task. The UI provides hierarchical control over the experiment. High-level task control allows the researcher to start, stop, or override or advance the experiment and adjust parameters like reward size. Finer-grained stimulation control enables precise, real-time adjustment of optogenetic parameters, including the stimulation epoch (e.g., inter-trial, choice) and the properties of light delivery (e.g.: pulse duration, frequency, and amplitude). Simultaneously, the interface offers comprehensive real-time feedback on animal performance. A status panel displays key metrics such as trial counts and choice rates, while a detailed history is plotted dynamically on a trial-by-trial basis. This allows researchers to immediately monitor learning progress, detect issues, and adjust protocols if necessary. In this example, the dashboard also integrates multi-modal feedback from peripheral hardware. Multi-camera views provide simultaneous video streams of the animal, while lick sensors connected to the choice spouts are displayed as binary indicators reflecting the current sensor status and provide a high-resolution sensor history centered on the choice interval.

Importantly, live-analysis can be performed alongside a task, for example, for quantizing multi-trial behavior for experiment decisions. Peripherals support dynamic interaction, either through parallel processing threads (e.g., real-time pose estimation) or via direct live plotting of activity, such as instantaneous readouts and time-stamped lick traces. Together, this interface enables comprehensive real-time observation and intervention, with the capacity to transform the dashboard into a centralized behavioral command center.

### Demonstrating Versatility in Rodent Behavioral Paradigms

To demonstrate Neurokraken’s real-world utility, we implemented a series of decision-making paradigms in mice (Figure 4). These examples showcase how the framework flexibly coordinates and integrates multiple off-the shelf and specialized hardware components and leverages a modular design to bridge distinct experimental domains. It further highlights the diversity in custom task logic flows based on a diverse range of input and output contingencies that can readily be implemented within Neurokraken. Alongside we present novel hardware components such as holders, motor mounts, and head-fixation towers, that were designed and 3D-printed, supporting rapid and flexible adaptation of the experimental hardware (see Supplementary Figure 1 and Supplementary Table 2, online documentation for designs).

**Figure 4:**
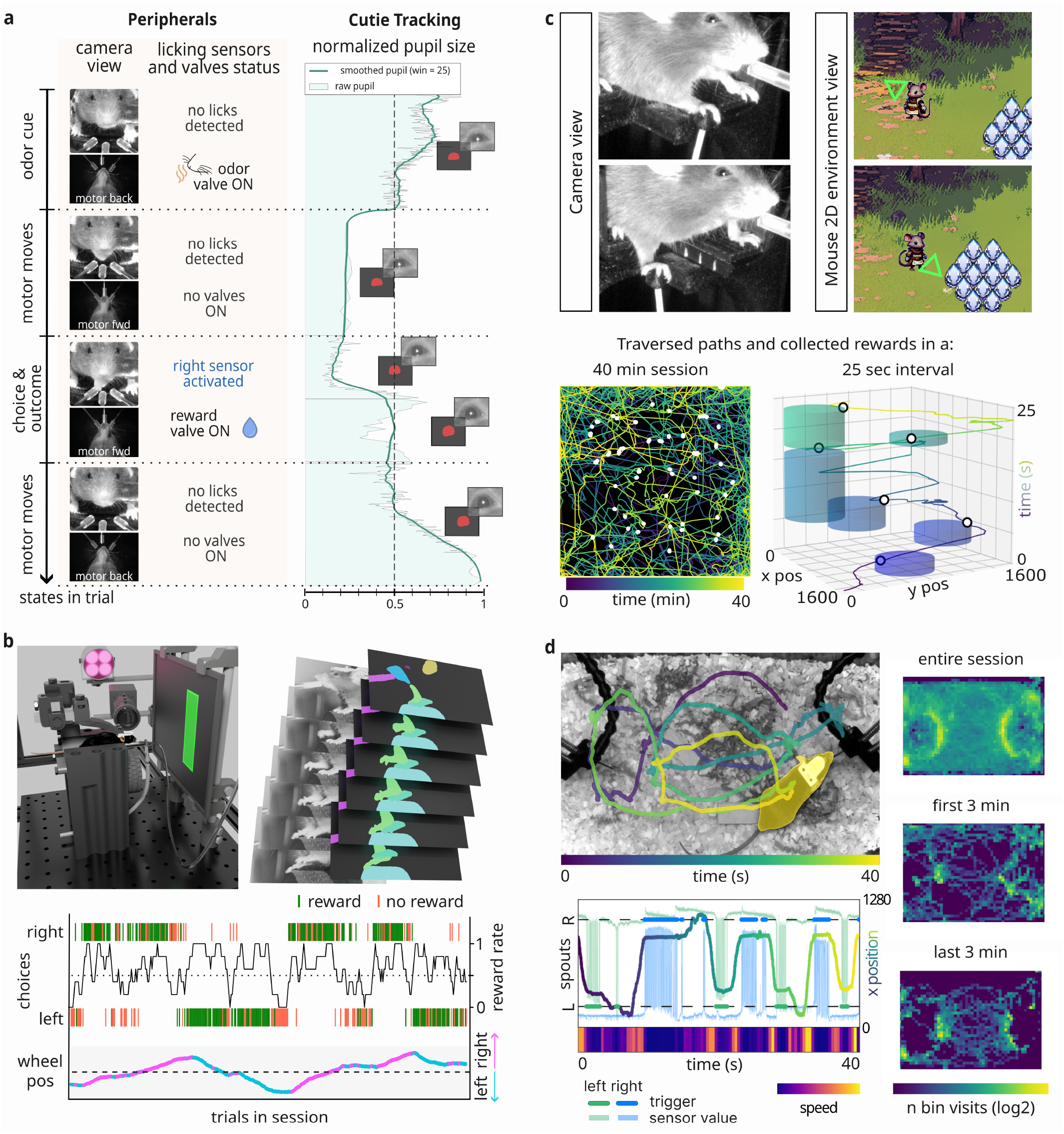
Versatile Implementation of Rodent Behavioral Paradigms. Neurokraken’s modular design capabilities demonstrated across four distinct paradigms, progressing from tightly controlled head-fixed tasks to freely moving, naturalistic environments **(a)** In an odor-guided bandit task, head-fixed mice perform a discrimination task using Neurokraken’s documented open olfactometer and 3 motorized choice spouts. The task states progress through odor presentation, spout movement, reading a choice from spout lick sensors, and potentially triggering reward delivery. The CUTIE object segmentation tool tracks pupil size. **(b)** In a modular adaptation, the lick sensors are replaced with a rotary encoder-connected steering wheel and the stimulus odors with a display. The mouse learned to steer a 2D object on a screen to the side providing a higher reward probability, providing a continuous readout of reaction time, movement angle, speed, and tracked body parts to quantify decision variables. **(c)** To study more complex spatial navigation, memory and cognition, a mouse navigates a 2D virtual arena. This setup allows for detailed analysis of exploration strategies, path optimization, and learning in dynamic environments. **(d)** Extending the framework to naturalistic behavior, mice interact with two spouts where licking one triggers a reward and trigger receptivity at the other, encouraging traversal. A top-mounted camera provides continuous trajectory tracking, visualized here as color-coded path histories and occupancy heatmaps, to quantify learning and other spontaneous behaviors over long timescales.

We first implemented a head-fixed olfactory discrimination task, leveraging the mouse’s innate reliance on smell (Fig. 4a). Using a custom-built olfactometer (see documentation and supplementary table 2 for further details), we delivered precisely timed odorants to the animal. Choices were made using three independently-motorized lick-choice spouts. The motorized movement provides both whisker and auditory feedback when a choice becomes available, and the spouts retracted instantaneously after incorrect choices, reinforcing learning through tightly coupled action-outcome associations. Licks were registered with millisecond precision by capacitive touch sensors (or, alternatively, infrared beam-break sensors for electrophysiology rigs), and correct choices triggered sucrose droplet delivery from the same spouts via solenoid valves. In parallel, Neurokraken captured continuous video from multiple camera angles, allowing for the live-analysis of, for example, pupil dynamics with the CUTIE model as a non-invasive proxy for attention and surprise^26^. Other python-available Computer Vision or AI tools can exchangeable be used live as part of the experiment or for post-experiment analysis of the saved video files^13,27^.

Highlighting the framework’s modularity, we then reconfigured the setup for a visuomotor control task (Fig. 4b). The lick spouts were replaced with a rotary encoder-connected steering wheel, which the mouse used to control a 2D object on a screen. By steering the shape to the left or right side on the screen, animals had to adapt to probabilistic reward changes associated with both sides. While CUTIE tracked relevant body parts, the encoder provided precise, continuous measurements of movement angle changes. These complementary readouts offer a powerful means of quantifying decision variables, such as reaction time, in dynamic environmental settings.

To investigate more complex cognitive processes, we designed a 2D virtual reality navigation task (Fig. 4c). Here, an avatar on the screen mirrored the animal’s movements, allowing it to explore a dynamically updating reward arena. This task exemplifies Neurokraken’s capabilities regarding the integration of unconventional devices into experiments, supporting highly interactive game-like continuous task designs, and adapting task states to corresponding custom visuals. This paradigm moves beyond simple binary choices, enabling the study of spatial learning, exploration, and adaptation under conditions that more closely approximate naturalistic uncertainty and can be as readily used by a human participant via a keyboard or other input device. Neurokraken tasks are not limited to 2D environments and can be designed and traversed in a 3D environment (see Figure 1b and online documentation).

To demonstrate Neurokraken’s capabilities beyond head-fixed preparations, we performed a freely moving, automated shuttling task in the home-cage (Fig. 4d). In this paradigm, mice learn that licking one spout triggers delivery of a sucrose-water reward and the readiness for the next reward at an opposing spout, encouraging repeated crossings of the cage. Throughout the task, frames from a top mounted camera were used to live-track the animals’ trajectories, revealing how rapidly they learn the task contingency and optimize their paths accordingly. In this example we quantified learning, but it allows as well tracking of a wide spectrum of naturalistic behaviors and ethological readouts, including social interactions, sleep patterns, and grooming within their home-cages.

#### Modular, open, 3D-printable hardware components

The Neurokraken platform is complemented by a library of versatile, 3D-printable components for constructing custom experimental setups (Supplementary Figure 1 and online documentation; models are freely available on printables.com). This collection includes connectors designed to mount hardware such as cameras, lights, and displays onto standard aluminum tubing. These parts provide full freedom for customization, increase availability if a cheap 3D printer is available, and reduce the cost of setups significantly due to the independence of commercial suppliers (Supplementary Figure 1a). 3D-printable breadboard baseplates with threaded inserts allow custom and firm placement of components in a modular approach according to experiment requirements. 3D-printable X, XY, and XYZ stages (Supplementary Figure 1b) further support adjustable fine-positioning of elements. For experiments utilizing head-fixed mice, we include an accommodating and simple to use cassette-based mounting solution that helps to reduce animal stress during head-fixation (Supplementary Figure 1c). Additionally, the library provides all 3D -printable parts and assembly instructions for a custom-built olfactometer, enabling odor-based behavioral paradigms. Neurokraken’s documentation further contains a walkthrough for building a behavior setup, and our table of cost-effective and useful components utilized for different neuroscience setups for an easy start or for scaling of experiments.

## Methods

### Neurokraken framework

The Neurokraken framework is a modular platform combining a high-level Python control layer with a Teensy 4.1 microcontroller for precise, low-level hardware management. Detailed instructions, example tasks, API references, feature guides, device wiring diagrams, and a complete setup assembly walkthrough are available on the official documentation site (https://Neurokraken.github.io/Neurokraken-book). The following sections summarize the key features and architectural principles relevant to the experiments described.

### Setup and Hardware Configuration

At the start of an experiment, all hardware components - such as sensors, actuators, cameras, microphones, and display settings are defined using Neurokraken’s *Configurator functions* (*configurators*.*py)*. These helper *functions* return standardized parameter dictionaries, providing an easy way to assemble a task setup while ensuring easy extensibility for researchers interested in adding new Arduino devices.

Configurators are used to instantiate the main *neurokraken* object. Sensors and actuators are provided as a single dictionary with user-defined key names (e.g., *‘reward_valve’*). After initialization, these keys are used within the *get* API object to read from sensors and send commands to actuators. The *neurokraken* object can also receive optional arguments, such as the log path or a flag to enable simulation/keyboard mode. As such, their primary job is to act as a user-friendly “builder” for hardware configuration. A researcher simply calls a *Configurator* function (e.g., *devices*.*rotary_encoder()*) and passes simple arguments, like the microcontroller pin numbers. *Configurators provide* standardized Python dictionaries containing all parameters that Neurokraken needs to understand and control that specific piece of hardware. This approach enables easy visibility, organization and configuration of peripherals.

A key feature of this process is the automatic generation of Arduino microcontroller code. A provided utility script (*config2teensy*.*py*) can be provided with a task’s code to automatically generate the required Arduino-side header file (*config*.*h*) from the connected devices. This enables the automatic generation of the hardware-side code, ensuring the Python logic and the microcontroller firmware are synchronized without the user ever needing to write C++ or Arduino code. This approach is central to Neurokraken’s “Arduino-free” philosophy. Thus, even advanced tasks can be fully defined within a single compact python script, as shown in our example documentation. Upon initialization, Neurokraken populates the global *get* (*neurokraken*.*controls*.*get*), making components accessible for all subsequent experiment code.

### Experiment Logic Definition

The behavioral logic is defined as one or more state objects. A state can contain a combination of 4 optional methods which define the core experimental logic. These methods are *loop_main()* which runs at kHz frequency dependent on the experimental setup (Supplementary Table 1), *on_start()* and *on_end() which can be used to define task behavior upon entering and progressing from a state, and loop_visual() to render custom graphics to a display*.

Live sensor and log data can be accessed via the *get* object and used to direct the progression of the state. While *loop_main()* executes with high temporal precision for time-sensitive logic, the *loop_visual()* speed is limited by the display’s refresh rate.

To support common state-machine paradigms, multiple states and their potential successors can be defined when loading the *experiment*. Within a state’s *loop_main()*, the code can set an outcome - such as progressing to a *‘reward’* or *‘timeout’* state based on the subject’s choice. This approach supports complex, branching logic while remaining simple to implement. Importantly, for continuous tasks, a single, persistent state can be used.

### Execution and User Interface Integration

Following the setup and logic definition, the task is started by calling the *neurokraken*.*run()* method. A key feature of this workflow is the seamless integration of user interfaces. A UI is implemented as standard Python code that is executed before the *neurokraken*.*run()* command. In this pre-run phase, UI elements (e.g., buttons, sliders) are defined and typically linked to callback functions which are executed upon interaction with the UI elements. Since *get* serves as the main interface to Neurokraken, it can also be used outside of the task logic from lines executed by UI elements to manually control task variables or connected devices. Similarly, *get* can access logged and live experiment data for UI elements to monitor experiment progression, behavior variables, update live-plots, or display live-camera feeds. This design of parallel execution ensures that the UI does not interfere with the core task’s timing while providing complete interactive control. We exemplify the usage of different popular python GUIs in our examples. Notably, usage of UIs (task displays) is optional and Neurokraken can be run without requiring a display for suitable tasks.

### Data Synchronization (Networking)

Communication between the Python host and the Teensy-based microcontroller is managed by the Networker component. It uses a high-speed request-response loop: Python sends a packet of control commands to be enacted, and the Teensy responds with a packet of new sensor data. To ensure data fidelity, the Teensy side maintains timestamped arrays of all unsent sensor changes with millisecond precision. These complete arrays are communicated to Python during each cycle for automated logging. This approach ensures that all sensor data is captured at millisecond precision, even if the host computer encounters processing delays, thus preventing data loss. Importantly, for devices requiring the highest temporal precision (e.g. the pulse clock and the multi-purpose “timed_on”) timing is implemented directly on the microcontroller through a function dedicated to the device’s highest framerate processes.

### Core Architecture and Deployment Modes

Neurokraken’s Python codebase consists of an internal *core* folder (managing the *networker, state_machine, main_loops*, etc.) and user-facing importable modules (*configurators*.*py, controls*.*py*). The framework can be deployed in two distinct modes: i) Imported Mode: This is the primary, flexible method described above, where Neurokraken is imported as a package into any Python script. This mode is ideal for flexible task development and custom-scripted experiments. ii) Runner Mode (*kraken*.*bat*): A command-line utility for high-throughput, standardized sessions. This mode operates on a task folder containing separate *config*.*py, task*.*py*, and optional files for UIs, AIs and subject files. The mode can pass common arguments (e.g., ‘--keyboard’ to run in keyboard mode or’-- config’ to use a specified file as *config*.*py*) and prompt for session metadata.

### Software and Dependencies

Experiments were validated on Windows with Python 3.13 (see online documentation for up-to-date information). Core dependencies include py5, pyserial, opencv-python, sounddevice and keyboard with some functionalities and demonstrated examples utilizing additional packages like harvesters, cutie, imageiio[ffmpeg], and ViZDoom. Installation is supported through conda, local virtual environments or uv (a python package manager), with step-by-step instructions provided in the documentation. The Arduino IDE (v2.2.1) firmware was compiled for the Teensy 4.1 using the Teensyduino board manager (1.59.0).

### Visual Environment Creation

The Neurokraken framework supports the integration of both 2D and 3D graphics, allowing for a wide range of visual complexity. Simple 2D graphical assets as used in our examples, such as icons and environmental textures, were generated using various open-source image generation models. These assets were then integrated into tasks using the Neurokraken’s loop_visual() which utilizes the py5 library for an extensive range of visual and animation capabilities, which is fully compatible with Neurokraken’s Python-native architecture. A Custom 3D corridor world was modeled and textured in Blender to exemplify the creation and integration of custom 3D virtual environments into Neurokraken tasks.

### Hardware and 3D-printed components

Sensors (e.g., beam breaks, lick/touch sensors), rotary encoders, valves, speakers, LEDs, motors, and small displays were sourced from common online suppliers (e.g., Amazon, Aliexpress, fluid concepts). Our online documentation’s hardware section maintains a list of accessible and effective parts suitable for neuroscience behavior setups, including store links, alternatives and price while the device library provides wiring guides to connect devices to the central teensy microcontroller coordinating sensors and actuators, time-stamping reading and synchronizing of external devices (https://Neurokraken.github.io/Neurokraken-book).

Structural components of the behavioral boxes and custom 3D-printed designs to accommodate peripheral components (examples shown in Figure 4) were designed in Blender 4.0 and exported for filament printing. Filament -based 3D printing was performed on Ender 3 V3 SE and V3 KE printers using PLA+ (eSun) or PLA (ecoPLA, 3DJake). See Supplementary Table 2 for details.

### System Performance and Latency Validation

We set up 5 tasks, with increasing compute demand reflected in up to 32 connected interacting devices and the addition of high framerate video capture and live-CUTIE processing. 10-minute-long hardware setup and input simulation tests were performed on a Windows 11 desktop workstation equipped with an Intel i7-11700K CPU and an NVIDIA RTX A4000 GPU. Video input was provided by a Kayeton USB camera operating at 720p and 90fps, with frames processed by CUTIE in its default configuration. Because the Networker transmits sensor data only upon detecting state changes, realistic load testing requires continuous input variability. We simulated high-frequency subject interactions using an external ESP32-S3 (QT Py) to drive digital inputs and servo motors, which in turn physically modulated rotary encoders. Analog signal variability was generated using photoresistors. To simulate output load, the Python main loop triggered random state changes in servo motors and valves throughout the session. We logged the precise timestamps of every main loop iteration and hardware communication cycle. To assess the integrity and completeness of sensor data we added a millisecond and microsecond sensor device to function as sensors whose values are certain to be different every millisecond, verifying that the system captures sensor data at millisecond precision without loss, even under increasing computational demand. To investigate the sub-millisecond sampling precision, the microseconds clock was added as the last sampled device where we expect the highest accumulating sampling time variation. By analyzing the logged microsecond at sampling time, we found that even in our most extreme test sensor sampling precision remained in the low microseconds (StdDev 1.08µs, Max Absolute 14µs). To validate the stability of the external time-alignment signal, a 100ms pulse clock was configured for each task. The output pin was connected to a secondary Teensy 4.1 microcontroller acting as an external recording device, which logged the arrival timestamps of the pulse edges at microsecond precision.

System performance was quantified by calculating the standard deviation (jitter) and absolute maximum latency of sequential timepoint differences for three key metrics: sensor value updates, main loop iterations, and clock signal pulses. This approach served to validate the Networker’s ability to ensure automated, millisecond-precise data collection independent of Python-side processing speed, and to confirm the reliability of the pulse clock as a ground truth signal for synchronizing external devices. All benchmark tasks and associated testing utilities are available in the *performance_test* directory of the Neurokraken online repository.

### Animal Experiments

C57BL/6J mice were group-housed (up to 5 mice per cage) in a temperature- and humidity-controlled facility under a 12-hour light-dark cycle. Except during water restriction for behavioral testing, mice had ad libitum access to food and water. In total 6 animals were used for the respective experiments. All animal procedures were approved by the local Animal Ethics committee of the Medical University Innsbruck and under the licenses GZ 2022.0.039.060 and GZ 2022-0.118.110 of the respective Federal Ministry of Women, Science and Research of the Republic of Austria.

## Discussion

Neuroscience currently demands the use of increasingly complex and multimodal behavioral paradigms that integrate advanced neural recording, manipulation, and computational analysis techniques. Historically, meeting this demand has been hampered by the high cost and technical rigidity of commercial systems, coupled with the fragmented nature of many open-source solutions. Our platform Neurokraken is designed to overcome these challenges by retaining laboratory-grade temporal precision while leveraging the flexibility of the standard Python ecosystem. This architecture delegates high-precision hardware control and synchronization to a microcontroller, while defining complex task logic entirely in high-level Python code. In combination, it provides the four essentials that define next-generation behavioral platforms: openness, flexibility, versatility, and cost-effectiveness. Particularly, the development focuses on versatility and flexibility stemmed from an effort to provide an easy-to-use software architecture to individuals new or relatively new to designing, building or adopting behavioral tasks.

Neurokraken is also designed for flexible integration of devices laboratories already own or can easily source common electronic parts from the most popular online stores accessible worldwide (e.g. Amazon or AliExpress). This further allows tasks to be replicated and implemented without difficulties in sourcing specific supplier-locked devices.

### Neurokraken Enables Translational and Complex Behavioral Design

Alongside the investigation to understand how the brain generates behavioral representations across species, a significant objective in brain research is to improve translational validity of rodent behavioral paradigms^28^. For example, to advance the understanding of neuropsychiatric disorders, while keeping ethology in mind, tasks must fractionate behavior into measurable and translatable motifs related to neuronal circuits that show at least a certain perseverance across species. Recently, new rodent tasks integrate or compare via human task parameters ^29,30^ for higher translational validity, which likely will benefit the predictive validity of neuronal perturbation experiments, whether through pharmacological or non-pharmacological means. Neurokraken is uniquely suited to support this paradigm shift, enabling the rapid construction and modification of highly controlled, complex cognitive tasks. Similarly, neuroscience is increasingly focused on computational neuroethology, the measurement and quantification of complex, naturalistic, and high-dimensional behaviors to understand brain function^1,31^. This requires seamless integration of real-time computational analysis with experimental control.

Neurokraken supports these demands through its multi-threaded architecture. This design ensures that computationally intensive processes, such as real-time pose estimation using deep learning tools like CUTIE^26^ or other suitable Python-integrable tracking solutions, can run in parallel threads. All these external processes can access live experimental data (like video frames or sensor readings) and influence the task progression through a unified mechanism (the *get* API). This capability supports behavior-driven closed-loop intervention, allowing features derived from movement or posture to be used for instantaneous feedback, thereby accelerating the integration of computational ethology into controlled systems neuroscience. The platform’s architectural separation, combined with Python’s versatility, also allows for the easy development of rich visual and virtual environments using libraries like py5, supporting tasks from psychophysics to complex virtual 3D navigation.

Neurokraken contributes to the growing open-source ecosystem, which includes seminal projects like BPod^32^, pyControl^20^, Rigbox^33^, Autopilot^34^, and specialized head-fixed systems like OHRBETS^24^ and HERBs^35^. While systems like pyControl focus on robustness and reproducibility via microcontroller firmware, Neurokraken with its many arms aims to complement and interconnect existing efforts by prioritizing software flexibility and access to the Python ecosystem that provides a significantly richer state-of-the-art development environment than MATLAB or other programming languages. This strict adherence to the actively maintained, standard Python ecosystem protects the platform against software decay, ensuring long-term community sustainability compared to tools built on insular codebases.

Neurokraken further addresses two major challenges prevalent in systems neuroscience: reproducibility and cost. The platform promotes reproducibility by providing a comprehensive repository of validated example tasks, high-level design logic, hardware integrability and configurability. It facilitates cost reduction by enabling researchers to construct highly precise rigs using easily sourced off-the-shelf electronic components and a library of versatile, cheap 3D-printed parts. This contrasts with standardized setups that often rely on expensive proprietary or difficult-to-procure components, particularly critical for labs with limited market access. Each of our behavioral task examples is test-ready with a budget of a few hundred euros. The successful, standardized approach demonstrated by collaborations like the International Brain Laboratory^12,16^ highlights the value of standardized protocols, hardware, and data analysis pipelines for achieving cross-lab reproducibility. Neurokraken was designed with this in mind and provides the architectural flexibility necessary to adapt and standardize complex paradigms in a similar, collaborative spirit.

### Limitations and Future Directions

Despite its versatility, Neurokraken also has inherent limitations. While the Python-native approach provides flexibility and despite our efforts to lower access barriers with examples and easy-to-use scripts and functions, users should possess some fundamental Python knowledge for advanced task customization. Fortunately, the recent rise in LLM-based coding assistance for Python has further reduced this barrier. Although Neurokraken is currently likely among the easiest open-source solutions for beginners as hardware configuration and integration is simplified, researchers completely new to electronics may still face challenges during initial setup, necessitating expanded documentation and the potential for pre-assembled hardware kits. As a fully open-source project hosted on GitHub, Neurokraken welcomes community contributions for users sharing their task or tool integration examples.

## Conclusion

Neurokraken offers a single, unified framework that merges hardware precision with software flexibility. By defining task logic in standard Python, the platform simultaneously solves the precision and complexity challenge in two transformative ways to shape the future of behavioral neuroscience: i) Deep Integration of the Scientific Python Ecosystem: The multi-threaded architecture and unified data access enables the seamless incorporation of sophisticated Python-based tools (including machine learning, psychophysics, and computational models) directly into the behavioral loop. This allows for real-time, behavior-driven experimentation and complex adaptive math models, thereby accelerating the critical shift toward computational neuroethology. ii)Cross-Species and Cross-Setup Task Reusability: The centralized, high-level task definition allows for task reusability across subjects and hardware setups. The same cognitive task logic can be deployed on diverse platforms - from an AI agent using environments like ViZDoom, to a head-fixed mouse performing complex sensorimotor control, to a human participant performing psychophysics on a touchscreen. This capability is helpful for translating cognitive paradigms across species and modalities, facilitating the comparison of neural computations and behavioral domains across the full spectrum of biological and artificial intelligence.

By merging laboratory-grade temporal precision with the flexibility of the modern Python ecosystem, Neurokraken empowers researchers to implement, share, and reproduce complex experiments at a fraction of the traditional cost. We hope this platform will catalyze the development of new adaptations and inspire and push further the continuing culture of openness and collaboration in behavioral systems research.

## Data and Code Availability

The data and code supporting the current study is openly available on https://github.com/Neurokraken/Neurokraken.

## Acknowledgements

We thank Maja Überegger for help with the animal work and Aron Koszeghy for sharing underlying data related to Figure 1h and Figure 4b. We further thank the many individuals behind open-source tools that enable Neurokraken and its features.

## Author contribution

All authors (A.W., S.C.A., J.P.) have contributed to the conception and design of the work. The codebase was written by AW. A.W., S.C.A., have performed the experiments, analyzed and visualized the data and provided the underlying code. A.W., S.C.A., and J.P. have drafted, revised and contributed to the article and approved the final version of the manuscript.

