## Supplementary Information for "Neurokraken: A fully flexible, open-source, python-based neuroscience behavior platform"

**Supplementary Material**

This Supplementary file provides additional materials supporting the main manuscript. It includes the modular 3D-printed hardware components used in the tasks shown in Figure 4 (Supplementary Figure 1); a table summarizing performance tests of Neurokraken under varying computational loads (Supplementary Table 1); and a representative parts list for Neurokraken-based behavioral tasks illustrated in Figure 4 (Supplementary Table 2).

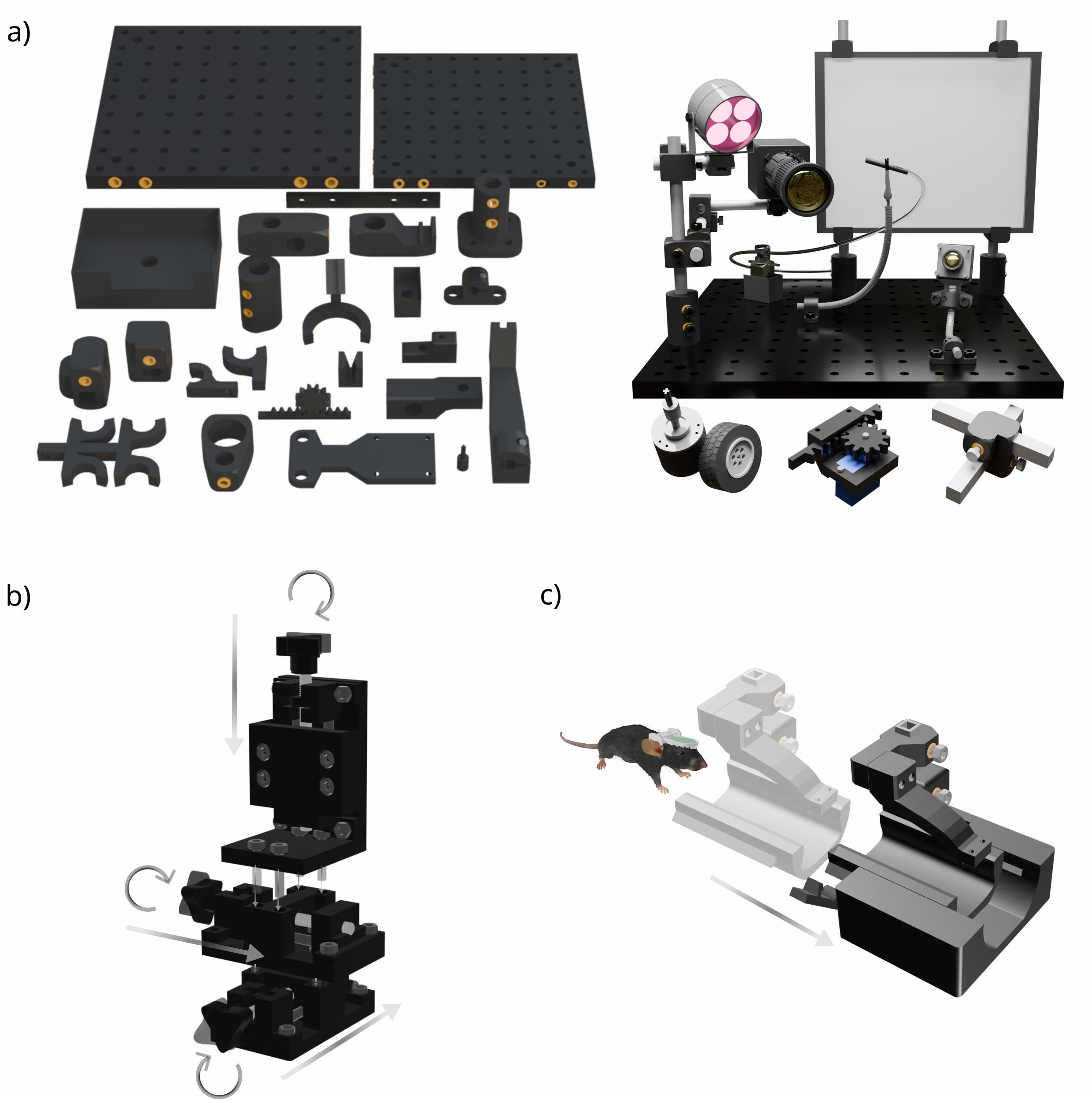

***Supplementary Figure 1: Modular 3D printed hardware* components developed for head-fixed Neurokraken-based mouse decision-making tasks. a)** connector and mounting components. **Left**: Selection of custom 3D-printed connectors, mounts, and holders used throughout the Neurokraken framework. These include multi-sized breadboards that can be screwed together in different configurations, several hardware holders, square and round rod connectors, a linear pinion-gear assembly, and syringe, spout and camera holders. ***Right**:* Example behavioral setup illustrating how these components can interface with optical breadboards and support some core hardware elements used in the decision-making paradigms shown in Figure 4, including cameras, displays, sensors, valves, motors, and a wheel coupler. **b)** Modular linear stages: 3D-printed linear positioning modules can be combined for a three-axis (X/Y/Z) stage enabling precise and independent adjustment of task components. **c)** Slide-in head-fixation cassette and station for head-fixed behavioral experiments.

**Supplementary Table 1: Performance test of Neurokraken under varying computational loads**

| | Performance Test Condition | | | | |
|---|---|---|---|---|---|
| | 1 rotatary encoder, 1 valve | 1 rotatary encoders, 1 valve, 90fps camera | 2 valves, 2 digital inputs, camera, Live CUTIE tracking | 10 digital inputs 5 servo motors | 10 digital inputs, 5 analog inputs, 2 rotary encoders, 10 valves, 5 servo motors |
| **Sensor data sampling** | 1ms | 1ms | 1ms | 1ms | 1ms |
| **Sensor sampling precision** | | | | | |
| STDEV | 0.30us | 0.27us | 0.33us | 0.66us | 1.08us |
| Maximum | 4us | 6us | 8us | 8us | 14us |
| **Python main loop** | | | | | |
| mean framerate | 2741.43it/s | 2012.05it/s | 1441.11it/s | 1957.63it/s, | 1430.87it/s |
| interframe STDEV | 0.48ms | 0.61ms | 0.68ms | 0.51ms | 0.52ms |
| interframe Maximum | 4ms | 9ms | 11ms | 7ms | 5ms |
| **Networker communication** | | | | | |
| mean framerate | 2741.24it/s | 2010.99it/s | 1439.81it/s | 1957.50it/s | 1423.19it/s |
| interframe STDEV | 0.48ms | 0.61ms | 0.69ms | 0.51ms | 0.52ms |
| interframe Maximum | 4ms | 9ms | 11ms | 7ms | 5ms |
| **Pulse clock synchronization on the receiver side** | | | | | |
| individual pulse time deviations STDEV | 0.90us | 0.92us | 0.90us | 1.38us | 62.39us |
| individual pulse time deviations Maximum | 7.20us | 6.80us | 7.10us | 10.21us | 109.21us |

**Supplementary Table 2: Representative parts list for Neurokraken-based behavioral tasks. H**ardware components, olfactometer, 3D-printing supplies, and other auxiliary components used across the decision-making paradigms in Figure 4. For each item the table provides the purchase link, supplier, price and the corresponding task(s) in which the component was used. Note prices will vary depending on region, tax and date and are understood to be indicative.

| TASK IN FIG 4 | ITEM AND LINK | SUPPLIER | PRICE [€] |
|---|---|---|---|
| All Tasks | **Microcontroller**<br>Teensy 4.1 | Exp Tech | 28.59 |

| | | | |
|---|---|---|---|
| | Aluminium heatsink for Teensy | Amazon | 4.88 |
| | **Sensors** | | |
| Task B | Rotary encoder | Amazon | 13.64 |
| Task A | Sparkfun capacitive touch sensor | Amazon | 6.68 |
| Task A | Optical interrupter photoelectric | Amazon | 5.87 |
| | **Cameras** | | |
| Task A & D | Mini camera Board | Amazon | 9.99 |
| All Tasks | 2.8 - 12mm Kayeton camera | Kayeton | 55 |
| | **Displays** | | |
| Task B & C | LP097QX1 IPS Panel with VSDISPLAY Controller | Amazon | 63 |
| | **Valves** | | |
| Task A | *6V Air Valve - FA0520E, ADA4663 - olfactometer* | Amazon | 8.98 |
| All Tasks | solenoid valve - 200010993 | Fluid Concept | 49.6 |
| All Tasks | Silicon Tubing 1.5mmx3.0mm 25m for solenoid valve | CARL ROTH | 25.1 |
| All Tasks | Mosfet module - Valve Controller | Amazon | 6.21 |
| | **Other Hardware** | | |
| Task B | Wheel 43.2mmx18mm. prod. Nr. 86652/32019 | Bricklink | 7.9 |
| All Tasks | IR light for cameras | Amazon | 15 |
| Task A | Step-down adjustable voltage converter | Amazon | 6.29 |
| All Tasks | Reward Spouts | Amazon | 6.85 |
| All Tasks – Electrophysiology setups | Alternative PTFE reward spots | Amazon | 8.00 |
| Task A | Servo motors | Amazon | 15.62 |
| Task A | **Olfactometer** | | |
| | Air filter regulator | Amazon | 14.9 |
| | Flow meter (0.1-1.5mL/min) | Amazon | 17.05 |
| | 5x6mm Air flow control | Amazon | 11.5 |
| | Manifold 5-ports 1/4-28 peek | Darwin Microfluidics | 217 |
| | Syringe filters (1-2um pore size; 15mm membrane) | CARL ROTH | 74.55 |
| | PTFE teflon tubing | PTFE Tube Shop | 7/meter |
| | PC4-M6 PTFE teflon tubes | Amazon | 9.24 |
| | T-connector pneumatic 6mm | Amazon | 10 |
| | Pneumatic straight reducer (4-6mm) | Amazon | 7.39 |
| | Stainless steel male thread M6 push | Amazon | 9.82 |
| | Push connection connectors M5 Male x 6mm | Amazon | 6.17 |
| | Luer connectors | Amazon | 6 |
| | Mineral oil | Amazon | 11.72 |
| | (R)-(+)-Limonene (Odor) | Sigma/Merck | 23.4 |
| | Hexanoic acid (odor) | Sigma/Merck | 21.9 |
| | Eugenol | Sigma/Merck | 105 |
| | Sabinene | Sigma/Merck | 134 |
| | trans-Cinnamaldehyde | Sigma/Merck | 77 |

| | | | |
|---|---|---|---|
| | D-Carvone | Sigma/Merck | 162 |
| All Tasks | **3D print**<br>Ender 3 V3 SE<br>Filament - ecoPLA 1.75mm<br>Filament - PLA+ Black eSUN, 1.75mm | <br>3DJake<br>3DJake<br>3DJake | <br>170<br>19.99<br>18.99 |
| All Tasks<br>To build Behavioral<br>Boxes | **Structural Components**<br>Threaded inserts M2-M6<br>Pack standard V-groove aluminium extrusion profile<br>Acrylic glass panels<br>corner brackets L shape<br>T-slot nuts M5<br>stainless steel rods used for linear stages<br>1cm aluminium square rod<br>1cm aluminium round rod | <br>Amazon<br>Amazon<br>Amazon<br>Amazon<br>Amazon<br>Amazon<br>Local Wholesale Würth GmbH<br>Local Wholesale Würth GmbH | <br>9.24<br>58<br>12.6<br>8.49<br>15.12<br>8.39<br>~5<br>~5 |
